# Cluster efficient pangenome graph construction with nf-core/pangenome

**DOI:** 10.1101/2024.05.13.593871

**Authors:** Simon Heumos, Michael F. Heuer, Friederike Hanssen, Lukas Heumos, Andrea Guarracino, Peter Heringer, Philipp Ehmele, Pjotr Prins, Erik Garrison, Sven Nahnsen

## Abstract

**Motivation:** Pangenome graphs offer a comprehensive way of capturing genomic variability across multiple genomes. However, current construction methods often introduce biases, excluding complex sequences or relying on references. The PanGenome Graph Builder (PGGB) addresses these issues. To date, though, there is no state-of-the-art pipeline allowing for easy deployment, efficient and dynamic use of available resources, and scalable usage at the same time.

**Results:** To overcome these limitations, we present *nf-core/pangenome*, a reference-unbiased approach implemented in Nextflow following nf-core’s best practices. Leveraging biocontainers ensures portability and seamless deployment in HPC environments. Unlike PGGB, nf-core/pangenome distributes alignments across cluster nodes, enabling scalability. Demonstrating its efficiency, we constructed pangenome graphs for 1000 human chromosome 19 haplotypes and 2146 *E. coli* sequences, achieving a two to threefold speedup compared to PGGB without increasing greenhouse gas emissions.

**Availability:** nf-core/pangenome is released under the MIT open-source license, available on GitHub and Zenodo, with documentation accessible at https://nf-co.re/pangenome/1.1.2/docs/usage.

## 1 Introduction

The availability of high-quality collections of population-wide whole-genome assemblies (Liao *et al*., 2023; Kang *et al*., 2023; Weller *et al*., 2023; Zhou *et al*., 2022; Liu *et al*., 2020; Leonard *et al*., 2022) offers new opportunities to study sequence evolution and variation within and between genomic populations. A challenge is simultaneously representing and analyzing hundreds to thousands of genomes at a gigabase scale.

One solution here is a pangenome. It models a population’s entire set of genomic sequences (Ballouz *et al*., 2019). In contrast to reference-based genomic approaches, which relate sequences to a linear genome, pangenomics relates each new sequence to all the others represented in the pangenome (The Computational Pan-Genomics Consortium, 2016; Eizenga *et al*., 2020; Sherman and Salzberg, 2020) minimizing reference-bias. Pangenomes can be described as sequence graphs which store DNA sequences in nodes with edges connecting the nodes as they occur in the individual sequences (Hein, 1989). Genomes are encoded as paths traversing the nodes (Garrison *et al*., 2018).

Current pangenome graph construction methods exclude complex sequences or are reference-biased (Chin *et al*., 2023; Minkin *et al*., 2016). One recent approach that overcomes such limitations is the PanGenome Graph Builder (PGGB) pipeline (Garrison *et al*., 2023). PGGB iteratively refines an all-to-all whole-genome alignment graph that lets us explore sequence conservation and variation, infer phylogeny, and identify recombination events. PGGB was already extensively evaluated (Garrison *et al*., 2023; Andreace *et al*., 2023) and applied to build the first draft human pangenome reference (Liao *et al*., 2023). However, PGGB is implemented in bash: This (a) makes it difficult to deploy on HPC systems, (b) does not allow for a fine granular tuning of computing resources for different steps of the pipeline (Sztuka *et al*., 2024), and (c) limits its cluster scalability because PGGB can only use the resources of one node. These limitations greatly hinder the broad application of large-scale pangenomes.

To compensate for that, we wrote *nf-core/pangenome*, a reference-unbiased approach to construct pangenome graphs. Mirroring PGGB, nf-core/pangenome is implemented in Nextflow (Di Tommaso *et al*., 2017). In contrast to PGGB, nf-core/pangenome can distribute the quadratic all-to-all base-level alignments across nodes of a cluster by splitting the approximate alignments into problems of equal size. We benchmarked the time spent on base-pair level alignments and show that it is reduced linearly with an increase in alignment problem chunks. We showcase the workflow’s scalability by applying it to 1000 chromosome 19 human haplotypes, and to 2146 *E. coli* sequences, which were built in less than half the time PGGB required while not increasing the CO2 equivalent (CO2e) emissions.

## 2 Material and Methods

### 2.1 Pipeline overview

The pipeline’s (Fig. 1a) input is a FASTA file compressed with *bgzip* (Li *et al*., 2009) containing the sequences to create the graph. Sequence names should follow the Pangenome Sequence Naming specification (PanSN-spec) (Garrison, 2021). The primary output is a pangenome variation graph (Garrison *et al*., 2018) in the Graphical Fragment Assembly (GFA) format version 1 (GFA Working Group, 2016).

**Fig. 1.**
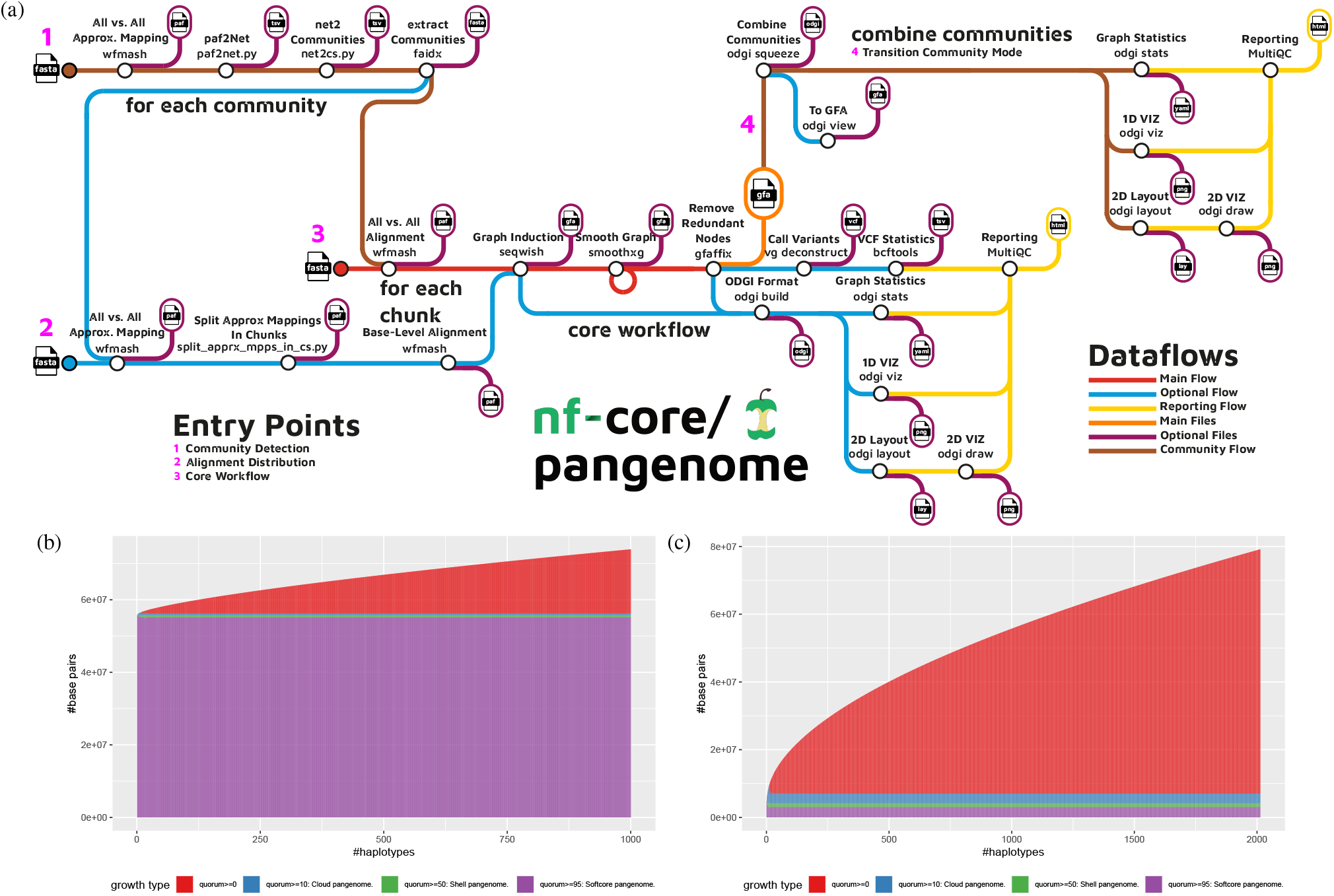
(a) Schematic representation of the nf-core/pangenome workflow processes and detailed analysis steps. The input consists of one FASTA file containing all sequences. The pipeline comes with 3 major entry points: Community detection (1), alignment distribution (2), and core workflow (3). Optional community detection (1) is performed on the input sequences. If selected, the heavy all-to-all baise-pair level alignments (2) can be split into problems of equal size. nf-core/pangenome’s core workflow (3) is a direct mirror of PGGB. If running in community mode, all communal graphs are combined into one (4) and the subsequent quality control subworkflow is executed. The output is a pangenome graph in GFA format. (b) + (c) Pangenome growth curves of the built pangenome graphs. Growth type is defined as the minimum fraction of haplotypes that must share a graph feature after each time a haplotype is added to the growth histograph. *quorum >*= 0: All sequences without any filtering are considered. *quorum >*= 10: Sequences traversed by at least 10% of the haplotypes. *quorum >*= 50: Sequences traversed by at least 50% of haplotypes. *quorum >*= 95: Sequences traversed by 95% of haplotypes. (b) Pangenome growth curve of the chromosome 19 pangenome graph of 1000 haplotypes. (c) Pangenome growth curve of the E. coli pangenome graph of 2013 haplotypes.

#### 2.1.1 Core workflow

The core workflow of nf-core/pangenome is an exact mirror of PGGB (Fig. 1a). The pipeline comes with additional enhancements: (a) All concurrent processes can be run in parallel. (b) Each process can be given individual computing resources.

The first step in the nf-core/pangenome pipeline is the all-to-all alignment of the input sequences with the whole-chromosome pairwise sequence aligner WFMASH (Guarracino *et al*., 2024). This avoids reference, order, or orientation bias, and allows each sequence in the pangenome to serve as a reference when exploring related variation. In the pangenome graph induction step SEQWISH (Garrison and Guarracino, 2022), an alignment to variation graph inducer, converts the sequence alignments into a variation graph. We then normalize the graph with the variation graph simplification algorithm SMOOTHXG (Garrison *et al*., 2023): A 1-dimensional (1D) graph embedding (Heumos *et al*., 2023) orders the graphs’ nodes to best-match the nucleotide distances of the genomic paths of the graph. Next, the graph is split into partially overlapping segments. The sequences of each segment are realigned with a local Multiple Sequence Alignment (MSA) kernel, partial order alignment (POA) (Lee *et al*., 2002). Afterwards, the segments are laced back together into a variation graph. By default, the SMOOTHXG process is applied 3 times in order to smoothen the edge effects at the boundaries of the segments. Finally, we employ GFAFFIX (Liao *et al*., 2023) to systematically condense redundant nodes within the graph.

Basic graph build quality is evaluated with ODGI: Optimized Dynamic Genome/Graph Implementation (Guarracino *et al*., 2022) for understanding pangenome graphs. ODGI reports basic graph statistics and diagnostic 1D and 2D visualizations. Optionally, nf-core/pangenome calls variants against any (reference) path(s) in the graph using *vg deconstruct* (Garrison *et al*., 2018). Finally, graph statistics and visualizations are summarized in a MultiQC (Ewels *et al*., 2016) report. Pipeline implementation details are given in Suppl. 5.1.

## 3 Results

### 3.1 Alignment jobs distribution evaluation

Generating all-vs-all alignments is a computationally quadratic problem. To evaluate nf-core/pangenome’s alignment jobs scalability (detailed in Suppl. 5.5), we applied it to 1024 *E. coli* genomes with varying numbers of chunks. nf-core/pangenome’s alignment jobs distribution linearly reduces the time spent on base-pair level alignments with increased chunks. The CO2 consumption is not influenced by the number of chunks (Suppl. Fig. 5.5).

### 3.2 Building a 1000 haplotypes chr19 pangenome graph

The Human Pangenome Resource Consortium (HPRC) recently built a draft human pangenome reference of 90 haplotypes (Liao *et al*., 2023). However, haplotype data for thousands of individuals already exists generated by the 1000 Genomes Project (1KGP) (Durbin *et al*., 2010). As a use case study, we used nf-core/pangenome to build a pangenome graph of 1000 chromosome 19 haplotypes (Kuhnle *et al*., 2020) within 3 days. The CO2e was 22.52 kg. PGGB built the same graph within 7 days. In Fig. 1b the pangenome growth curve generated with PANACUS (Liao *et al*., 2023) shows a growth of the number of nucleotides with an increasing number of haplotypes. The size of the softcore pangenome does not change with increasing numbers of haplotypes.

### 3.3 Building a 2146 sequences *E. coli* pangenome graph

To evaluate the pipeline’s scalability, we built a pangenome graph of 2146 *E. coli* sequences. nf-core/pangenome built the pangenome graph in 10 days, emitting 175.18 kg of CO2e. Due to wall clock time restrictions on our cluster, PGGB was not able to finish the graph construction within 30 days. To build a reasonable pangenome growth curve (Fig. 1c) we dropped all paths containing “plasmid” (130 in total) in their name. The softcore pangenome of the graph does not change with an increasing number of haplotypes (stable at 3Mb of sequence), but the general growth curve is steep.

## 4 Discussion

We implemented nf-core/pangenome, an easy-to-install, portable, and cluster-scalable pipeline for the unbiased construction of pangenome variation graphs. It is the first pangenomic nf-core pipeline enabling the comparative analysis of gigabase-scale pangenome datasets. The pipeline’s core workflow steps were already successfully applied to *Neisseria meningitidis* (Yang *et al*., 2023), wild grapes (Cochetel *et al*., 2023), humans (Guarracino *et al*., 2023; Liao *et al*., 2023), grapevines (Guo *et al*., 2024), taurines (Milia *et al*., 2024), and rats (Villani *et al*., 2024) underpinning the community effort to focus on a best-practice workflow to create reference-unbiased and sequence complete pangenome graphs. The modular domain-specific language (DSL) 2 pipeline structure eases the exchange of key processes with alternative tools, the extent of the pipeline with new tools, and the integration of parts of the pipeline with other (sub-)workflows.

We have shown that we are able to perform all-vs-all base pair level alignments of thousands of sequences. When executed on an HPC, nf-core/pangenome’s parallel workflow accelerates graph construction compared to PGGB. PGGB’s inability to assign individual computational resources to each pipeline step leads to the allocation of one whole node of an HPC, despite the fact that some processes can only make use of one thread. This blocks valuable CPU cycles for other users working on the same HPC and ultimately can lead to additional costs. In contrast, nf-core/pangenome does not have such limitations: Nextflow’s process management enables the optimal workload of given compute resources which can be especially important when running a pipeline in commercial clouds.

Competing pipelines don’t use any workflow management system to connect their processes (Chin *et al*., 2023), or their workflow language of choice is e.g. Toil (Vivian *et al*., 2017; Hickey *et al*., 2023) which makes them less user-friendly, less cluster efficient, and less portable (Wratten *et al*., 2021). nf-core/pangenome is currently the only pangenomics pipeline that is optionally monitoring its CO2 footprint. The measurements have shown that constructing extensive pangenome graphs, such as the 2146 *E. coli* graph, requires a considerable amount of energy. Therefore, before executing environmentally questionable experiments, we would recommend thoroughly assessing both the rationale and the methodology.

Although, we expect our pipeline to scale well for future pangenome graph construction challenges, such as for the next HPRC phase which targets 300 individuals, there still is potential for further optimization: IMplicit Pangenome Graph (IMPG) (https://github.com/ekg/impg), a tool that extracts homologous loci from all genomes mapped to a specific target region. This would allow us to break the whole genome multiple alignments into smaller pieces, construct a pangenome graph for each piece, and lace these together into a full graph with https://github.com/pangenome/gfalace.

We anticipate the pipeline, or its parts, will enhance current single linear reference analysis methods to explore whole population variation instead of focusing on one reference only. Looking ahead, pangenome construction pipelines like nf-core/pangenome will play a pivotal role in studying entire populations, single-cell whole genome sequencing analysis, and constructing personalized (medical) pangenome references (Sirén *et al*., 2023).

## Software and data availability

Code and links to data resources used to build this manuscript and its figures, can be found in the paper’s public repository: https://github.com/subwaystation/pangenome-paper.

## Acknowledgments

We thank Matthias Seybold from QBiC for maintaining the Core Facility Cluster. We thank Sabrina Krakau from QBiC for giving feedback to the nf-co2footprint plugin section. We are grateful to the nf-core community for their support during the implementation of the pipeline. From the nf-core community, we want to thank Matthias Hörtenhuber, Maxime Garcia, Susanne Jodoin, Julia Mir Petrol, Adam Talbot, and Gisela Gabernet.

## 5 Supplement

### 5.1 Implementation

nf-core/pangenome is written in Nextflow using its latest domain-specific language (DSL) 2 syntax which facilitates a modular pipeline structure. Each software tool is an individual process that is implemented in its own module (https://nf-co.re/docs/contributing/modules). Processes are concatenated into subworkflows (https://nf-co.re/docs/contributing/subworkflows). Developed with the nf-core framework, the pipeline follows a set of best-practice guidelines ensuring high-quality development, maintenance, and testing standards. Specifically, we provide community support via a dedicated Slack channel (https://nfcore.slack.com/channels/pangenome), GitHub issues, and detailed documentation (https://nf-co.re/pangenome/1.1.2/docs/usage). Versioning and portability are enabled through (a) semantic versioning (https://semver.org/) of the pipeline via tagged releases on GitHub, (b) packaging software dependencies in archivable containers so that the software compute environment is the same across different systems, and (c) summarizing software versions and parameters in the MultiQC report of the pipeline. nf-core/ pangenome uses biocontainers to facilitate portability across different computing resources like HPC clusters, cloud platforms, or local machines. Code changes are evaluated with GitHub Actions’ continuous integration (CI) using a pipeline-specific small test data set. For each new pipeline release, a full-size test is run on Amazon Web Services (AWS) validating the code integrity and cloud compatibility of real-world data sets. Specifically, a pangenome graph is created from the 8 Saccharomyces cerevisiae strains of the Yeast Population Reference Panel (YPRP) (Yue and Liti 2018). The results of such a run are available on the nf-core webpage (https://nf-co.re/pangenome/1.1.2/results/pangenome/results-0e8a38734ea3c0397f94416a0146a2972fe2db8b). Because we implemented our processes using DSL2 nf-core/modules (https://github.com/nf-core/modules), they can be distributed easily to other users to share commonly used processes or subworkflows across pipelines. This boosts the reuse of existing work done by the community to be integrated into future pipelines.

### 5.2 Chromosome community detection

Eukaryotic genomes are usually organized into chromosomes. Taking this into account during graph construction, the chromosome groupings from the input sequence are examined. Specifically, the homologies detected in the all-to-all WFMASH mapping step are put into the Leiden (Traag *et al*., 2019) clustering algorithm. The edge weight is *mapped*_*length* * *mapped*_*identity*. For each of the resulting communities, the nf-core/pangenome core workflow is executed in parallel. The communal graphs are joined into one and a final round of quality control is applied (see Fig 1a, brown tubes). In practice, this works well for large input sequences with a large mapping length (>1Mb) filter, which was demonstrated by Guarracino *et al*. (2023) when exploring the recombination between heterologous human acrocentric chromosomes.

### 5.3 Compute environment

We applied the nf-core/pangenome pipeline v1.1.2 to various inputs evaluating both the scalability of the all-vs-all alignment step as well as the pipeline as a whole. We used Nextflow version 23.10.1.5891 and Singularity version 3.8.7-1.el8 for each pipeline run. Experiments were conducted on our core facility cluster (CFC) with 24 Regular nodes (32 cores / 64 threads with two AMD EPYC 7343 processors with 512 GB RAM and 2 TB scratch space) and 4 HighMem nodes (64 cores / 128 threads with two AMD EPYC 7513 processors with 2048 GB RAM and 4TB scratch space). Each Nextflow process was given at most 64 threads. This ensures a fair run time comparison with PGGB v0.5.4 which was always executed on one Regular node via Slurm.

### 5.4 Estimation of the carbon footprint of pipeline runs

We also estimated the carbon dioxide equivalent (CO2e) emissions of each nf-core/pangenome pipeline run using the nf-co2footprint Nextflow plugin (https://github.com/nextflow-io/nf-co2footprint) v1.0.0-beta. Using the Nextflow resource usage metrics and information about the power consumption of the compute system, the plugin first estimates the energy consumption for each pipeline task. It then uses the consumed energy’s location-specific carbon intensity to estimate the respective CO2e emission. The calculations are based on the carbon footprint computation method developed in the Green Algorithms project (www.green-algorithms.org) (Lannelongue *et al*., 2021).

### 5.5 Alignment jobs distribution

The computationally heavy all versus all base-pair level alignments can be distributed across nodes of a cluster: First, WFMASH is run in mapping mode (WFMASH MAP), finding all sequence homologies using approximate alignments. The resulting Pairwise mApping Format (PAF) file is split into chunks of equal problem size. The number of chunks is manually selected. The value can be guided by the number and size of the input sequences, and by the available hardware. Assuming the number of chunks equals the number of nodes on a cluster, then potentially each base-pair level alignment (WFMASH ALIGN) can be run in parallel on each node (Fig. 1a, cyan tubes). All resulting PAFs are then forwarded to the pipeline’s core workflow which is continued at the SEQWISH process.

**Fig. S1.**
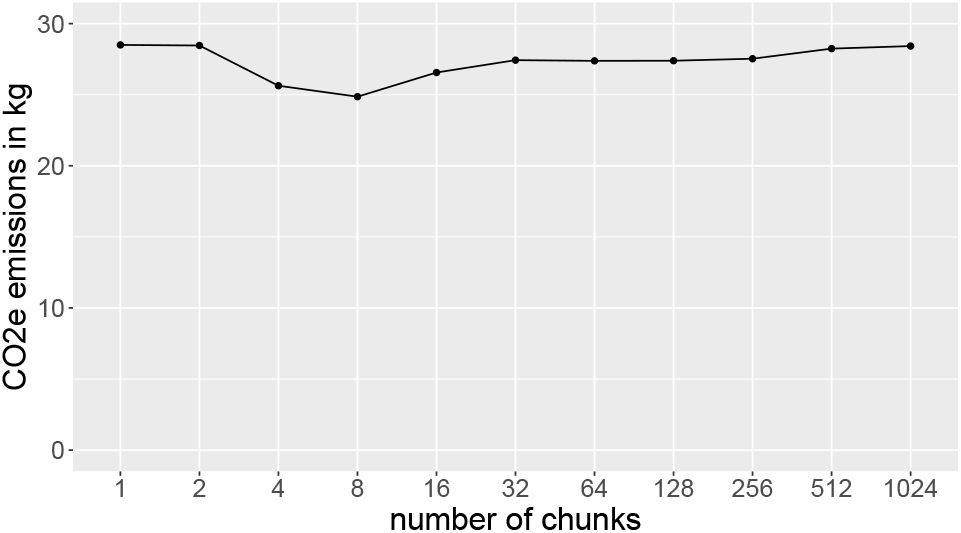
Base-pair level alignment evaluation. CO2e emissions are stable across varying numbers of chunks.

### 5.6 1KGP chromosome 19 data set

The FASTA of the chromosome 19 data set was downloaded in December 2023 from http://dolomit.cs.tu-dortmund.de/chr19.1000.fa.xz. The data set is described in (Kuhnle *et al*., 2020). Statistics of the built pangenome graph can be seen in Supplementary Fig. 5.6. The initial graph contains over 97% of Ns. We applied *odgi crush* (odgi version 0.8.6), which crushes consecutive Ns of all nodes containing Ns into just one N per node, to the 1KGP chromosome 19 pangenome graph. This brings down the number of Ns from over 3B to exactly 6000 (Suppl. Fig. 5.6). The 2D visualization (Suppl. Fig. 5.6) is perfectly linear without any large SVs present hinting that only short-read data was used to create the haplotype sequences. In contrast, the 2D layout of the HPRC PGGB chromosome 19 pangenome graph (Heumos *et al*., 2023) clearly presents SVs, especially in the centromere’s location. Investigating these complex regions of a human chromosome is only possible when using long-read assemblies for graph construction.

**Fig. S2.**
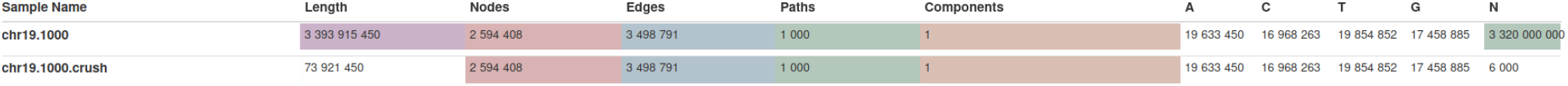
Screenshot of the output of ODGI’s MultiQC module displaying the vital graph statistics calculated by odgi stats of the 1000 Genomes Project 1000 haplotypes chromosome 19 pangenome graphs. In the crushed graph consecutive Ns of all nodes containing Ns were merged into just one N per node. A: Number of adenine bases in the graph. C: Number of cytosine bases in the graph. T: Number of thymine bases in the graph. G: Number of guanine bases in the graph. N: Number of bases with unknown base identity.

**Fig. S3.** odgi draw 2D layout displaying the graph topology of the crushed 1KGP pangenome graph. Structural variation would appear as bubbles.

### 5.7 *E*.*coli* data set

The 2146 full length E. coli sequences originate from Genbank (Sayers *et al*., 2021) and were downloaded 18 months ago. The initial pangenome consisted of 2 graphical components (Suppl. Fig. 5.7). This means that no strong homologies were found in some sequences. There can be many reasons for additional graph components: (a) The chosen sequence identity during the WFMASH mapping was not low enough. Although we went for a low 90% sequence identity (as was done by Garrison *et al*. (2023)), we still observe this additional graph component, so its sequence must be quite dissimilar to all other sequences. (b) There is human contamination in the bacterial sequences (Breitwieser *et al*., 2019). (c) Some sequences from GenBank may be of a low quality or were misassembled. We then used odgi explode to extract the largest graphical component, applied *odgi crush* and dropped all paths containing “plasmid” in their path name with *odgi paths*. This left us with one component and 2013 paths. In the 2D visualization, we observe a highly connected graph (Suppl. Fig. 5.7). All the reasons mentioned above, but especially horizontal gene transfer could explain this phenomenon. Therefore, there are a lot of edge crossings in the pangenome graph. The long stretches is dangling sequence. We speculate that here the 88 thousand Ns could play role.

**Fig. S4.**
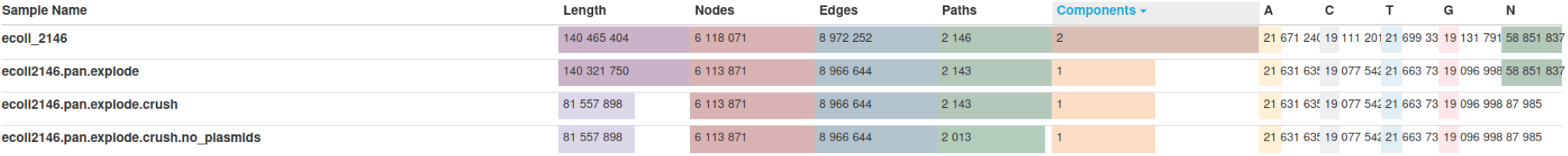
Excerpt of the 2146 sequences E. coli pangenome graph’s MultiQC report. Displayed are vital graph statistics by MultiQC’s ODGI module. A: Number of adenine bases in the graph. C: Number of cytosine bases in the graph. T: Number of thymine bases in the graph. G: Number of guanine bases in the graph. N: Number of bases with unknown base identity.

**Fig. S5.**
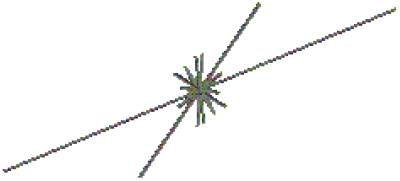
odgi draw 2D layout visualization of the 2013 haplotypes E. coli pangenome graph.

